# A canonical cortical electronic circuit for neuromorphic intelligence

**DOI:** 10.1101/2025.03.28.646019

**Authors:** Maryada, Chiara De Luca, Arianna Rubino, Chenxi Wen, Matteo Cartiglia, Ioan-Iustin Fodorut, Melika Payvand, Giacomo Indiveri

**Affiliations:** Institute of Neuroinformatics, University of Zurich and ETH Zurich, Zurich, Switzerland; Digital Society Initiative, University of Zurich, Zurich, Switzerland; Department of Information Technology and Electrical Engineering, ETH Zurich; IMEC, Leuven,Belgium; Astrali,Romania

## Abstract

Cortical microcircuits play a fundamental role in natural intelligence. While they inspired a wide range neural computation models and artificial intelligence algorithms, few attempts have been made to directly emulate them with an electronic computational substrate that uses the same physics of computation. Here we present a heterogeneous canonical microcircuit architecture compatible with analog neuromorphic electronic circuits that faithfully reproduce the properties of real synapses and neurons. The architecture comprises populations of interacting excitatory and inhibitory neurons, disinhibition pathways, and spike-driven multi-compartment dendritic learning mechanisms. By co-designing the computational model with its neuromorphic hardware implementation, we developed a neural processing system that can perform complex signal processing functions, learning, and classification tasks robustly and reliably, despite the inherent variability of the analog circuits, using ultra-low power energy consumption features comparable to those of their biological counterparts. We demonstrate how both the model architecture and its hardware implementation seamlessly capture the hallmarks of neural computation: attractor dynamics, adaptation, winner-take-all behavior, and resilience to variability, within a compact, low-power computing substrate. We validate the model’s learning performance both from the algorithmic perspective and with detailed electronic circuit simulation experiments and characterize its robustness to noise. Our results illustrate how local, biologically plausible rules for plasticity and gating can overcome challenges like catastrophic forgetting and parameter variability, enabling effective always-on adaptation. Beyond offering insights into the nature of computation in neural systems, our approach introduces a foundation for ultra-low power, fault-tolerant architectures capable of complex signal processing at the edge. By embracing -rather than mitigating-variability, these neuromorphic circuits exhibit a powerful synergy with emerging memory technologies, suggesting a new paradigm for sophisticated “in-memory” computing. Through such tight integration of neuroscience principles and analog circuit design, we pave the way toward a class of brain-inspired processors that can learn continuously and respond dynamically to real-world inputs.

## 1 Main

Neuromorphic processing systems are electronic systems designed to implement models of biological neural networks with various degrees of biophysical realism [1–3]. In this work we focus on systems built using analog Complementary Metal-Oxide-Semiconductor (CMOS) circuits that directly emulate the dynamic properties of biological neural processing systems through their physics [4, 5]. These circuits exhibit numerous similarities and share many constraints with biological neural circuits [6, 7]. The relationship between these circuits and neural processing systems “*is rooted in the shared limitations imposed by performing computation in similar physical media*.” [8]. Therefore, the use of analog neuromorphic technology represents “*a powerful medium of abstraction, suitable for understanding and expressing the function of real neural systems*.” [8]. The remarkable alignment between the physics of neuromorphic electronic circuits and the biophysical substrates of neural circuits [7, 9, 10] can be exploited on one hand for fundamental research, to validate or invalidate theories of neural computation, and on the other for applied research, to develop compact and power-efficient brain-inspired computing systems that directly exploit the physics of their computing substrate for realizing their operations.

A wide range of analog neuromorphic CMOS circuits have been proposed to directly emulate neural dynamics [10–14] and build mixed-signal Spiking Neural Network (SNN) chips [15–22], for both basic and applied research. More recently emerging memory technologies and novel memristive devices have also been proposed as non-volatile memory elements that can be used as analog neuromorphic circuit elements to directly reproduce many features of real synapses [23–29]. Computing systems built using these circuits and devices, similar to the biological architectures they emulate, typically carry out complex processing with extremely low energy figures, compared to conventional synchronous digital processors [6, 19, 30]. However, similar to their biological counterparts, they are affected by a high degree of variability, mainly due to device mismatch [31, 32].

Often considered a flaw or a “bug” in these circuits, variability and heterogeneity has been recently shown to be helpful (i.e., a “feature”) in theoretical models of neural computation for improving computing and coding performance [33, 34], robustness in learning [35], generalization in deep networks [36], and for enabling flexible computations involving multiple timescales [37–39]. Indeed, reliable and robust computation can be achieved in variable and noisy computing substrates, as in biology and analog neuromorphic/memristive systems, pursuing three main strategies, which are ubiquitous in biological brains: one is to use *populations* of synapses and neurons, connected via recurrent excitatory-inhibitory (E-I) pathways [33, 40, 41]; the second is the use of *disinhibition* as a gating mechanism (i.e., the suppression of inhibitory connections to enable the propagation of signals) [42–44]; and the third is the implementation of *learning* and adaptation [45–47]. In particular, recent learning models based on local spike-driven plasticity, dendritic compartments, and stop-learning criteria modulated by multiple factors, have been shown to be especially promising [34, 47–51].

Within this context we present a novel neuromorphic “canonical” cortical circuit motif [52, 53] that uses (i) a population of excitatory and inhibitory silicon neurons, in a balanced regime; (ii) a dis-inhibition pathway comprising multiple types of inhibitory cells; and (iii) a local spike-based dendritic learning mechanism, which has been *co-designed* to satisfy both theoretical neuroscience modeling and analog electronic circuit design considerations simultaneously, following a bottom up approach [54]. In particular, we show how computational neuroscience methods and principles of neural design [55] are instrumental in guiding the construction of efficient, robust, and accurate artificial neural processing systems, when the underlying computing substrate shares similarities and constraints with biology. For the analog neuromorphic computing substrate case, we use electronic neuron and synapse circuits designed with transistors operating in the *weak-inversion domain* [7]. They are ultra-low power leaky Integrate-and-Fire (I&F) neuron circuits with spike-frequency adaptation mechanisms that have biologically plausible time constants and are compatible with advanced Fully-Depleted Silicon on Insulator (FDSOI) 22 nm technology nodes [30]. The learning circuits exploit the dis-inhibition properties of the cortical circuit motif to seamlessly switch between training-mode and inference-mode, addressing the catastrophic forgetting problem, and endowing artificial sensory-processing systems the ability to provide always-on learning features on continuous data streams in real-world applications.

We demonstrate how the network proposed can achieve robust and reliable performance on signal processing and learning tasks, both with hardware-aware behavioral simulations and detailed circuit simulations.

While recent computational studies are starting to show the importance of variability in specific sets of parameters of spiking neural networks [33–35, 37–39] this is the first work that models in high-level hardware-aware simulations the effect of device mismatch of all of the network parameters, including weight values, spike threshold, time constants, adaptation rates, and refractory periods, validating the model with experiments with analog neuromorphic circuit-level simulations. In addition, we present for the first time local synaptic plasticity weight-update circuits that are compatible with multiple neural network learning rules, including simple Delta-rule perceptrons [56, 57], Bienenstock-Cooper-Munro (BMC) theory [58], Spike-Timing Dependent Plasticity (STDP) [59–61], and Behavioral Time Scale Synaptic Plasticity (BTSP) [62, 63].

We argue that neuromorphic processors built following this co-design approach can be used as fault-tolerant always-on in-memory-computing systems, ideally suited for edge-intelligence applications, able to operate reliably also when interfaced to emerging memory technologies such as nanoscale memristive devices [32]. By providing them with a powerful and robust local learning rule, this work paves the way to building a novel generation of ultra-low power brain-inspired adaptive signal processing technologies that can complement and outperform conventional computing ones in many specialized application areas, solving many of the challenges that standard digital technology is starting to face [64–66].

## 2 Results

By using heterogeneous populations of excitatory and inhibitory spiking neurons, the cortical circuit motif proposed minimizes the effects of variability thanks to both its recurrent excitatory and inhibitory connections and its dendritic learning rule.

### 2.1 The spike-based learning canonical microcircuit cortical motif

The diagram of the proposed circuit motif is depicted in Fig. 1a. It is a recurrent neural network that represents a “canonical microcircuit” [52, 53, 67] which can be configured to operate as an attractor network for modeling working memory [68], as a soft winner-take-all network for decision making [67], as an intrinsic oscillator [69, 70] or as an encoding layer for amplifying, filtering, and compressing high dynamic range input signals [71–73]. Furthermore, when learning is enabled, it can be trained to form robust attractors, classify signals into multiple classes, or recognize spatio-temporal patterns. The network relies on using a population of neurons connected in a excitatory-inhibitory balanced configuration for significantly reducing the effect of variability in its analog circuit implementation. The device mismatch that affects these analog circuits typically leads to Coefficient of Variations (CVs) of approximately 0.2 [10, 40]. In accordance with the Central Limit Theorem [74], a populations of uncoupled *N* silicon neurons can lead to a reduction of the standard deviation by a factor of 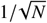. However, recent results in computational neuroscience have shown how variance can scale *superclassically* as 1/*N* in recurrent networks operating in balanced regimes [33, 75]. The network depicted in Fig. 1a exploits these results and extends their robustness features by endowing them with a local dendritic learning mechanism.

**Figure 1:**
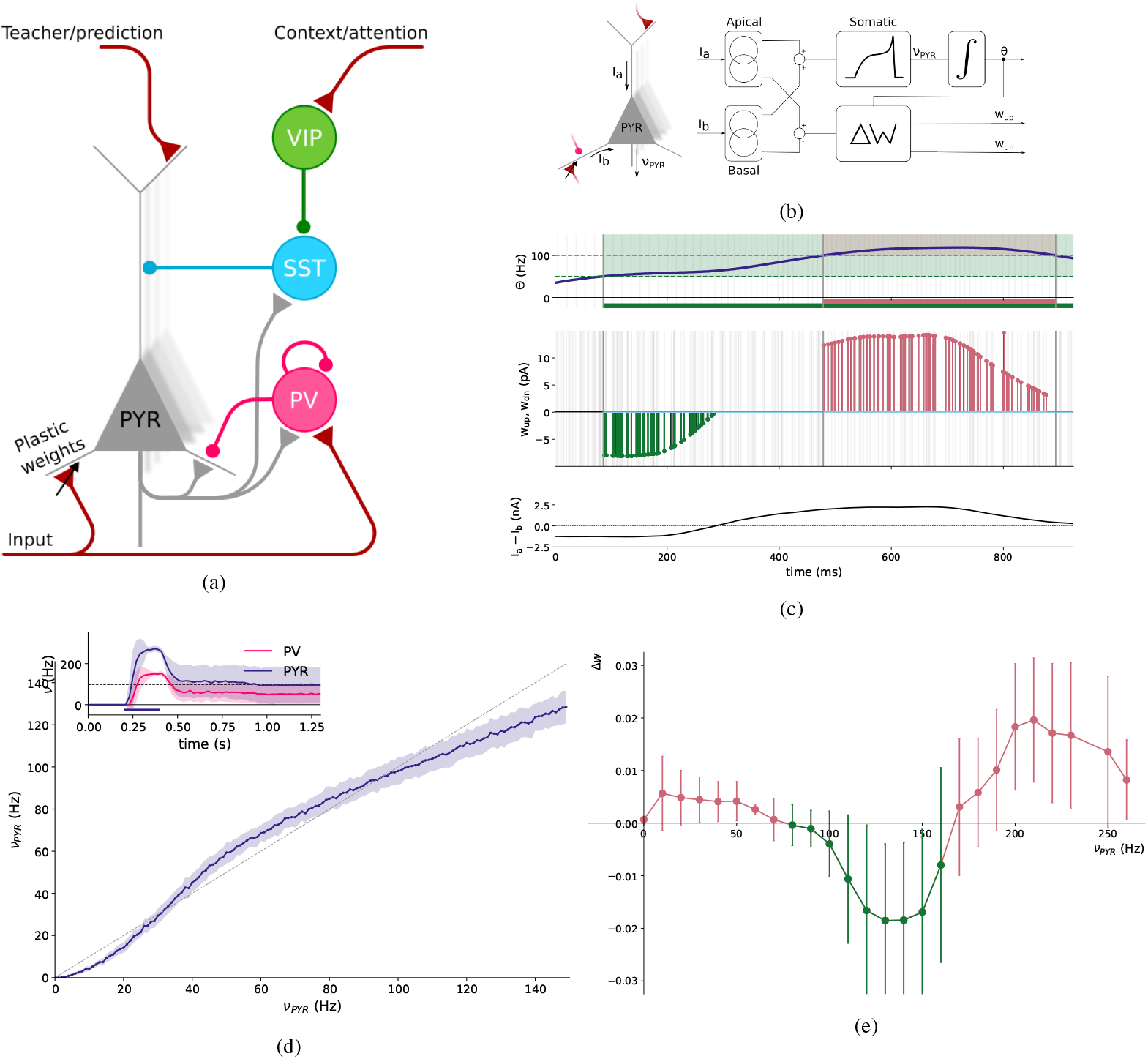
Canonical microcircuit motif with stable attractor and learning properties. (a) Circuit connectivity diagram. The “PYR” element represents a population of excitatory pyramidal neurons. The “PV”, “SST” and “VIP” elements represent the different populations of inhibitory neurons; (b) Block diagram of the “PYR” silicon neuron circuits used to integrate the basal and apical dendritic currents and to calculate the weight update control signals *w*_*up*_ and *w*_*dn*_; (c) Synaptic weight updates as a function of the post-synaptic neuron’s average firing rate (*θ*, top plot) and the error signal (*I*_*a*_ − *I*_*b*_, bottom plot). Weight changes (middle plot) are applied only upon the arrival of the pre-synaptic spike (shaded vertical lines). Positive/negative updates (*w*_*up*_/*w*_*dn*_) can only be applied if the *θ* variable lies within the corresponding LTP/LTD region (green and red areas in the top plot). The values of the currents on the ordinates of the plots are indicative, as the simulations could only use educated estimates for the scaling factors of the internal Very Large Scale Integration (VLSI) circuit currents; (d) Effective response function of the canonical microcircuit operating as an attractor network: the attractor stable fixed point is at about 20 Hz, consistent with the average firing rate plotted in the figure inset, after removal of an external driving input current (dark bar in the bottom part of the figure inset). (e) Rolling average of the weight update distribution versus the postsynaptic neuron firing rate during the training phase of a classification task, showing BMC-like dynamics with distinguishable LTD and LTP regions.

The network comprises one class of excitatory Pyramidal (PYR) neurons and three classes of inhibitory neurons: Parvalbumin (PV) cells, Somatostatin (SST)-expressing cells and Vasoactive Intestinal Peptide (VIP)-expressing ones. The pyramidal neurons have three compartments: a somatic compartment, a basal, and an apical synaptic input compartment. The somatic compartment models the Adaptive Exponential Integrate-and-Fire (AdExp-IF) neuromorphic circuit [10, 76], which can reproduce a wide variety of neuron dynamic behaviors [77]. The dendritic apical compartment implements a half-wave rectification non-linearity allowing currents to pass through only if they are larger than a set threshold (see Supplementary Material, Section SM 0.1). There are four different types of synapses with corresponding neuromorphic circuit implementations [13]: an excitatory one with slow *N*-Methyl-d-aspartate (NMDA) time-scales, an excitatory one with fast *α*-Amino-3-hydroxy-5-methyl-4-isoxazolepropionic Acid (AMPA) dynamics, an inhibitory *γ*-Aminobutanoic Acid (GABA) synapse circuit with subtracting effects (GABA_B_ type), and an additional inhibitory GABA circuit with both subtracting and shunting effects (GABA_A_ type).

All PYR neurons have recurrent excitatory connections and receive bottom-up sensory inputs to their basal compartment through plastic NMDA synapses and top-down ones through static AMPA synapses. The top-down inputs targeting the PYR neuron apical dendrite, represents a “teacher” signal in case of supervised learning frameworks, or a “prediction” or “prediction error” signal in case of predictive learning frameworks. The PV neurons implement both feed-forward and recurrent inhibition to establish E-I balance and soft Winner-Take-All (WTA) function. The SST neurons are excited by the PYR neurons and inhibited by the VIP ones. When active, they strongly inhibit (i.e., *gate*) all excitatory currents from the PYR neuron apical compartments to their soma compartments. The VIP neurons are stimulated by another class of top-down signals that represent an “attention” or “context” cue. When stimulated by these top-down signals, the VIP cells strongly suppress (dis-inhibit) the SST neurons, allowing the apical currents of the PYR neurons to pass through to their soma without any attenuation. This VIP-SST-pyramidal neuron dis-inhibitory mechanism has been shown to play important computational roles in a wide variety of modeling studies [42, 53]. It serves as an automatic way to enable/disable learning, controlled by top-down signals produced either by other cortical motifs in other parts of the network (e.g., in other layers of multi-layer networks), or by other external spiking inputs.

#### Attractor dynamics

Attractors are a hallmark of robust and stable neural computation [78], crucial for implementing stateful networks capable of pattern storage and recall [79]. It has already been shown that computational models of SNNs can form stable attractors [68, 80], and these theories have already been validated with mixed-signal analog/digital implementations on neuromorphic processors [81, 82]. More recently we have also shown how the properties and stable points of attractors can be set to desired values, by means of cross-homeostatic inhibitory plasticity both with theoretical models and neuromorphic processor validations [83]. Here we show that the cortical motif of Fig. 1a is also compatible with these theories and can form stable attractor states, despite the variability in all the network parameters, as affected by device mismatch present the analog circuit implementation. Figure 1d shows the effective response function of the canonical microcircuit. As shown, the recurrent excitatory PYR-PYR coupling combined with the negative feedback through the PYR-PV coupling gives rise to the classical response function exhibiting three fixed points, allowing sustained activation of ensembles at the non-null stable fixed-point, even after the input is removed.

#### The dendritic delta learning rule

The dendritic learning rule implemented in the model and by the corresponding neuromorphic circuits is driven by a three-factor [62, 84] spike-based Hebbian weight update mechanism derived from the original Delta-rule [56]: when a plastic synapse on the basal dendrite receives an input spike, its weight is updated proportionally to the difference between the total apical input current and the basal one, gated by a third-factor which depends on the level of the post-synaptic neuron’s Calcium concentration. This third-factor “stop-learning” gating signal enables (or disables) synaptic weight updates of the corresponding post-synaptic neurons if its past average activity proportionals to its Calcium concentration lies within (or outside) a set Long Term Potentiation (LTP) or Long Term Depression (LTD) learning region. This error-based, gated learning rule is crucial for providing a balance between plasticity and stability in a continuous learning framework [45, 85].

Figure 1b shows a block diagram of the learning circuits. The sum of all excitatory and inhibitory postsynaptic currents from the apical compartment are denoted as *I*_*a*_ and the ones from the basal compartment as *I*_*b*_. The “Apical” and “Basal” blocks in the figure implement current mirror circuits that sum copies of *I*_*a*_ and *I*_*b*_ into the somatic compartment block, and subtract them in the weight update learning block. Specifically, the learning block calculates an error signal *δ*(*t*) as:

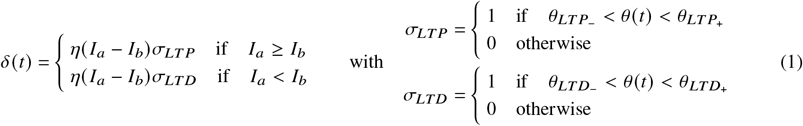

where *η* is a learning rate parameter, *θ*(*t*) is a proxy for the neuron’s Calcium concentration, calculated as the integral of the post-synaptic pyramidal neuron’s firing rate by a Differential Pair Integrator (DPI) current-mode circuit [13]. The variables 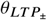 and 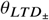 are the LTP and LTD upper and lower stop-learning thresholds, and *σ*_*LT P*_ and *σ*_*LTD*_ are the stop-learning gating variables for the LTP and LTD weight updates respectively. The detailed contributions to the basal current *I*_*a*_ and the apical one *I*_*b*_ are provided in Section 4.3.

While the error signal *δ (t*) is computed in continuous time and shared across all synapses of the same PYR neuron, the local weight update of an individual NMDA plastic synapse *i* is computed as: Δ*w*_*i*_ = *δ(t*_*i*)_, where *t*_*i*_ is the time of the arrival of the pre-synaptic spike at that synapse. Figure 1c shows an example of how the learning circuits update the weights of a synapse, as it is being stimulated by pre-synaptic input spikes, and as the post-synaptic mean activity *θ(t*) evolves over time. The top trace shows the temporal evolution of *θ(t*), computed by integrating the post-synaptic neuron output spikes (gray vertical lines). The two shaded regions mark the area where LTD and LTP are enabled (green and red respectively). The bottom plot shows the evolution of the error signal *δ(t*). The middle plot shows the negative and positive weight updates, applied with every pre-synaptic input spike (gray vertical lines). As prescribed by the theory, these jumps are proportional to the amplitude of the error signal and contingent on *θ*(*t*) falling within the respective learning regions (e.g., note that when the error *δ(t*) in the bottom plot turns positive at around *t* = 250*ms*, there are no weight updates in the middle plot, because *θ(t*) in the top plot lies in the LTD-only region).

In addition to being a local learning rule compatible with both the requirements of the analog neuromorphic circuits implementation and theoretical Calcium-based dendritic learning rules [50, 85], this rule is also consistent with the predictions of the BMC theory. Figure 1e shows the average weight changes during the training phase of a binary classification task (described in the next section), as a function of the post-synaptic neuron’s mean firing rate.

### 2.2 Binary classification of overlapping patterns

Here we demonstrate a binary classification task using two patterns made of binary vectors of independent Poisson spike trains, with either a high mean firing rate (50 Hz), or a low one (5 Hz) (see also Section SM 2.1). The binary classification network is implemented by splitting the neural populations of the circuit motif in two identical copies, with the only difference in the inputs to their apical dendrite (see “Teacher A/B” inputs in Fig. 2a). Figure 2b shows how the learning evolves as the network is trained to distinguish between the two input patterns, when they have with 50% overlap (see left box of Fig. 2d and Section SM 2.1). Specifically, we define an overlap *ov* between two input vectors of length *N* as the fraction of shared active synapses, where the number of active synapses for a given input is 0.5 × (1 + 0.5 * *ov*) × *N*. Since the number of active synapses varies with overlap, we are in a non-constant input current scenario, making the task more challenging for an SNN compared to maintaining a fixed number of active synapses while shifting the input. During training, a common “attention” signal is sent to all VIP cells of all columns to enable learning by dis-inhibiting the SST cells. This allows the PYR cells to fire at higher rates for entering their corresponding LTD and LTP regions (green and red shaded areas in Fig. 2b). As the input pattern is provided to all PYR neurons of all columns, a class-specific (“A” or “B”) high-teacher signal is provided only to the column of the corresponding class. Neurons that receives both the input and the high teacher signal are driven to bring their Calcium concentration, represented by the *θ* variable, in their LTP region. Conversely, neurons receiving only the basal input but no teacher apical inputs tend to fall in their LTD region, leading to depression of the stimulated synapses. After multiple (10 examples) presentations of the input and corresponding teacher signals, the weights of non-overlapping inputs converge on average to sufficiently high values for the true class inputs, and low values for the false class ones. The weights of the unspecific overlapping inputs are driven to intermediate values and do not contribute to the classification of the input patterns. Once the common “attention” signal is removed, the SST cells become active and reduce the average activity of the PYR neurons below LTP and LTD regions, ensuring an “inference” only stable operation mode (see SM 2.1.1 section for an analysis on the role of dis-inhibition mechanism). The performance of the network, for the 50% input overlap case, as a function of different levels of device mismatch variability is shown in Fig. 2c. Figure 2d shows the input patterns used, the weight matrices of the corresponding “A” and “B” columns, averaged over multiple training sessions, and the ideal weight matrix of a classifier not affected by variability. Although the change in synaptic weights is analog, their absolute value is discretized to 4 bit resolution, both in the software simulation and in the hardware implementation. The three center weight matrix plots show their initial randomly initialized state, their state at the end of the training, and after multiple “inference” repetitions, demonstrating that they remain stable once the network enters inference mode. The data in Fig. 2b and Fig. 2d was obtained for a device-mismatch variability factor of 20% as a “worst-case” scenario to account for corner cases in the chip fabrication.

**Figure 2:**
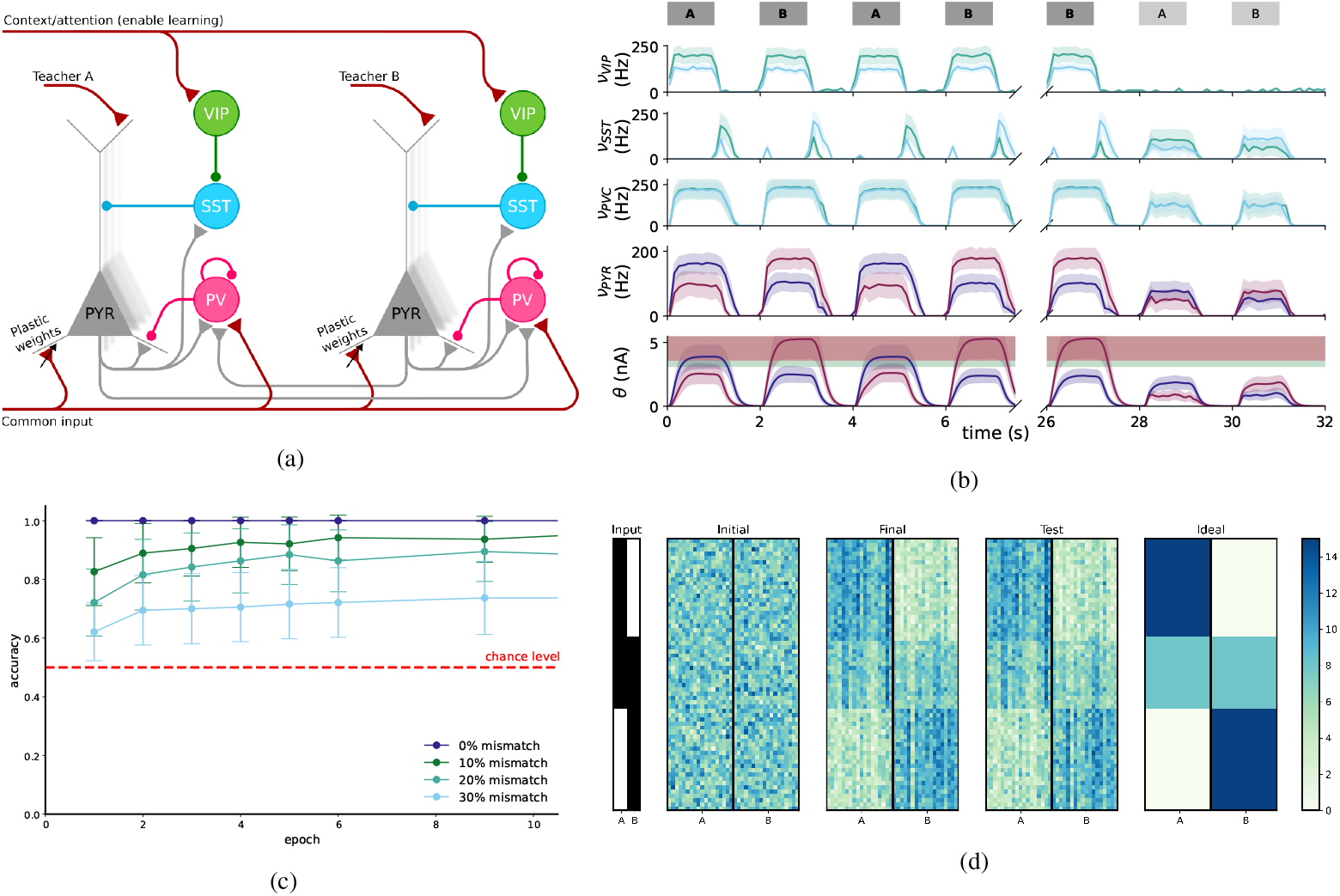
Binary classification experiment. (a) Network connectivity diagram with distinct teacher signals; (b) Population average firing rates during training and inference phases in response to different inputs with and without teacher signals. The *θ* variable in the bottom plot represents the time-averaged rates of the PYR neuron outputs. (c) Network performance for inputs with 50% overlap, as a function of simulated device mismatch variability; (d) Evolution of synaptic matrix weights matrices during training Evolution of synaptic weight matrices during training for a 50% overlap, meaning that more synapses are active so that the number of overlapping synapses equals % of the active synapses in the non-overlapping scenario.

### 2.3 Analog VLSI circuit implementation

The analog neuromorphic circuits used to implement the cortical motif of Fig. 1a comprise the ultra-low power silicon neuron, already presented in [30], the excitatory and inhibitory versions of the DPI silicon synapse [10, 13], as well as a novel Calcium-based dendritic learning circuits co-designed with the computational model proposed. The learning circuits are subdivided into two main parts: a novel circuit that computes in continuous time the error term *δ*(*t*) (eq. 1) in the neuron somatic block, and multiple copies of a novel circuit instantiated in the plastic (basal) input synapses which update their corresponding synapse weight upon the arrival of the pre-synaptic input spike, by an amount that depends on the error term computed in the soma.

Figure 3a shows the schematic diagrams of the soma-block error circuit. The circuit calculates the positive and negative parts of the error term *I*_*up*_ and *I*_*dn*_ by subtracting, in continuous time, copies of the basal *I*_*b*_ and apical *I*_*a*_ currents, within a tolerance level *I*_*ε*_, and multiplies this by a learning rate current *I*_*η*_, which is gated by the third factor *σ*_*LT P*_ and *σ*_*LTD*_ voltage variables, as defined in eq. (1). The synaptic weight update circuit is shown in Fig. 3b. This circuit produces a change in the synaptic weight of their corresponding plastic synapse by sampling the *I*_*up*_ and *I*_*dn*_ currents calculated in the soma, when its pre-synaptic input spike *V*_*s pk*_ is active. The charge produced by this sampling process is added or removed from the weight capacitor *C*_*w*_, increasing or decreasing the weight internal variable *V*_*w*_ accordingly. In parallel this voltage is slowly driven to the middle point between its maximum value (*i*.*e*., the power supply rail *V*_*dd*_) and its minimum value (*i*.*e*., ground) by a slew-rate limited amplifier in negative-feedback mode [46]. As this analog voltage reaches its power-rail limits, a standard digital 4 bit counter circuit then acts as a Analog-to-Digital Converter (ADC), discretizing and storing the weight value in Static Random Access Memory (SRAM). The combination of this stability mechanism on an analog voltage, with a 4 bit digital counter ensures robustness to noise [45, 86] and provides a number of discrete weight levels that has been shown to be sufficient for solving a wide range of network level benchmarks [87].

**Figure 3:**
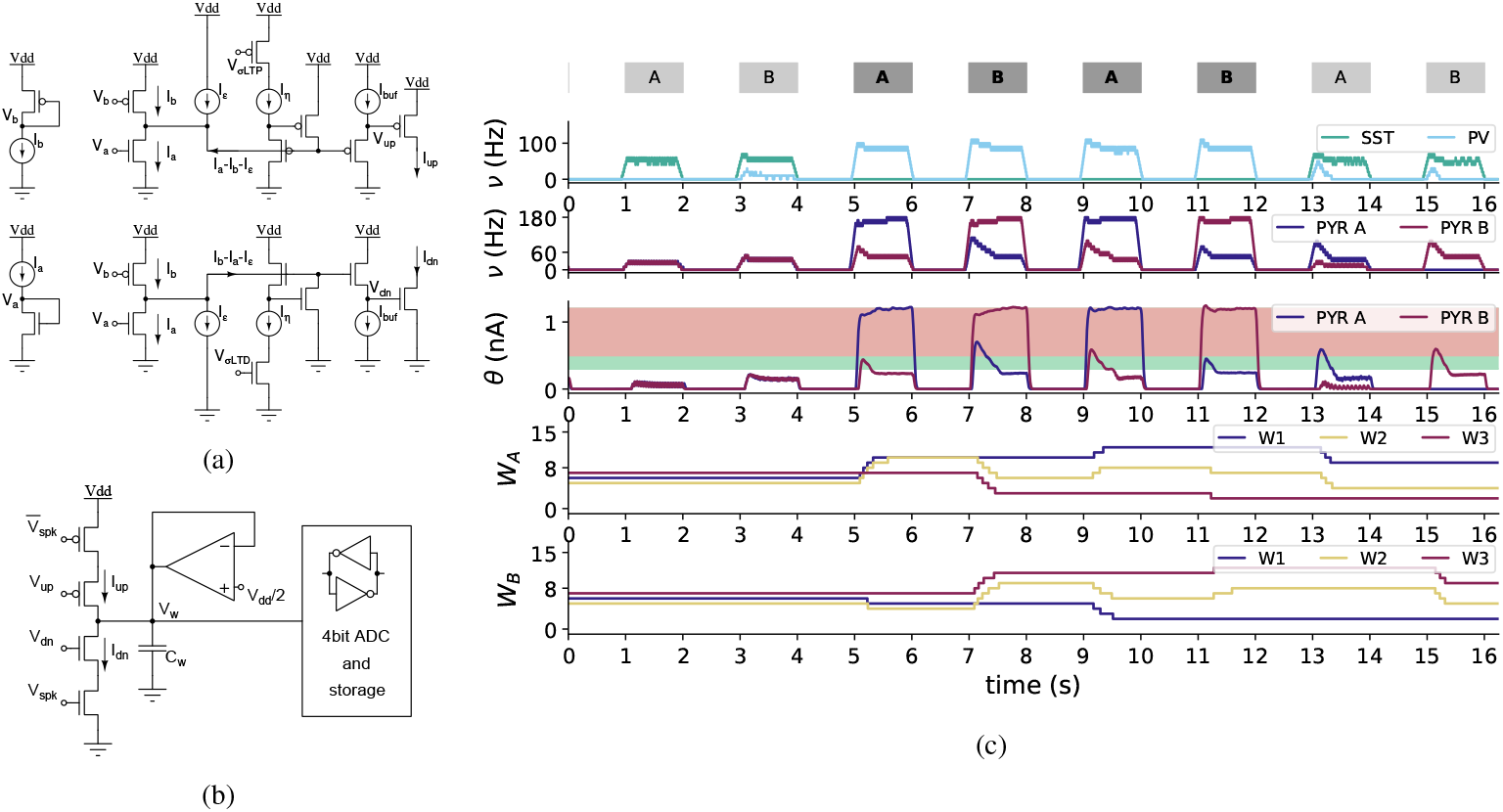
Learning circuits. (a) Soma-level error calculating circuits. (b) Synapse-level weight-update and storage circuit. (c) Circuit simulation results with overlapping patterns experiment.

Figure 3c shows circuit-level simulation results for the same binary classification experiment of overlapping patterns performed in Section 2.2. In this case, due to the extensive simulation times required by the circuit-level simulation tools, we only considered single neuron of each type (i.e., one pyramidal neuron per class, one inhibitory PV neuron, one SST and one VIP neurons). Each pyramidal neuron has three plastic synapses with 4 bit synaptic weights. As for the input pattern shown in Fig. 2d, the input pattern ‘A’ is formed by providing a high spike rate to synapses with weights W1 and W2, while pattern ‘B’ stimulates synapses with weights W2 and W3. The synapse with weight W2 therefore is uninformative, as it receives inputs from both patterns. At the beginning of the experiments, the plastic synaptic weights —W1, W2, and W3 —for both pyramidal neurons are initialized to the same random values, so that both respond in the same unspecific way to both input patterns ‘A’ and ‘B’ (see responses to the first two presentations of the input). After 5 seconds, training is turned on, and in addition to the input pattern, a high teacher signal is provided to the apical compartment of the neuron circuit of the corresponding class (i.e., pyramidal neuron of class ‘A’ receives both input and a high teacher when the pattern presented is ‘A’ and only input when the pattern is ‘B’). As the neuron’s Calcium concentration current, denoted as *θ(t*) enters the LTD and LTP regions, the weights are decreased or increased, depending on the sign of the error signal. As expected, for the neuron of class ‘A’, the weight of the first synapse increases, the weight of the third one decreases, and the weight of the middle synapse is driven to an intermediate value. For the neuron of class ‘B’, the third weight undergoes LTP, the first one LTD and the middle one is driven, on average, to the middle. After training, from 12 seconds on, the two neurons exhibit proper tuning to their corresponding input patterns.

## 3 Discussion

Neuromorphic computing has the potential of providing a revolutionary technology that can support ultra-low power always-on operation for processing low data-rate signals in edge computing applications. This is particularly true when the neuromorphic systems developed employ analog circuits operating in the weak inversion regime for signal processing, and/or memristive devices for non-volatile data storage. However, this technology has not been adopted at large so far because there are two main challenges to solve: on one hand, both analog circuits and memristive devices are affected by a high degree of variability, which hinder their reliability and reproducibility; on the other hand, the lack of standardized design methodologies and programming frameworks for spiking neural networks makes it difficult to efficiently develop and deploy these solutions at scale.

Adopting the same strategy pursued by Misha Mahowald and Carver Mead in their first neuromorphic engineering design efforts [6], this work showed how applying the principles of neural computation used by biological neural circuits to artificial spiking neural networks, co-designed and compatible with their electronic neuromorphic circuits, can solve both challenges at once. First we showed how the use of population of neural circuits, configured as recurrent E-I balanced networks can produce stable and reliable activities, even with large amounts of variability (e.g., with a CV of up to 20%). Then we showed how a biologically realistic dis-inhibitory cortical circuit motif can be used to naturally switch from training mode to inference mode, using a Hebbian spike-driven learning mechanism that only needs a top-down cue or context “label” signal for implementing a robust gradient-like delta rule.

We demonstrated the robustness of the cortical motif proposed, with both large network-level simulations that take into account the transistor-level non-idealities of their CMOS circuit implementations, as well as detailed CMOS circuit-level simulations. While the synapse and neuron CMOS circuits adopted have already been presented in the literature, we also presented novel CMOS circuits for implementing the local spike-driven learning rule that incorporates a Calcium-based learning region which enhances the robustness of the weight update mechanism, making it less susceptible to disruption from noisy sensory inputs.

A key feature of the hardware architecture lies in its ability to perform always-on -chip learning, which is essential for continuous adaptation, even when pre-trained networks are deployed. In contrast to many existing spike-based learning rules, typically validated only via bit-precise software simulations or digital spiking neural network simulators, our approach relies exclusively on locally available information at each neuron, both in space and in time [47]. By calculating the learning error continuously, in real-time, the architecture can robustly adapt to changes in both the internal signals and the external inputs, minimizing the effect of noise and variability.

The proposed mixed-signal electronic architecture and its on-chip learning via a calcium-based dendritic rule, represents a critical step in the evolution of adaptive, low-power neuromorphic systems. The co-design approach followed, by developing neural processing and learning rules hand-in-hand with the corresponding electronic circuits, validates expected behavior directly on silicon and provides a solution specifically tailored for mixed-signal neuromorphic hardware. Future efforts will extend this framework by incorporating homeostatic learning rules [83, 88] and integrating enhancements for temporal processing within a multi-layer topology, thereby further advancing the capabilities and scalability of neuromorphic systems.

## 4 Methods

### 4.1 The neuron model

The selected neuron model used is a multi-compartment AdExp-IF neuron in which currents from two dendritic branches, the bottom-up basal branch (*I*_*basal*_) and the top-down apical branch (*I*_*a pical*_), get integrated into a somatic compartment as membrane current (*I*_*soma*_):

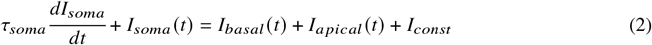

The basal and apical dendritic dynamics are given by:

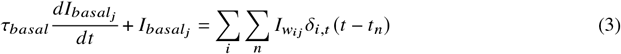

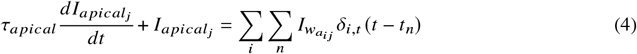

where *I*_*w*_ = *w* × *I*_*bias*_, where *I*_*bias*_ is the analog current representing the weights, and *w* is the 4-bit digital weight learned to scale *I*_*bias*_.

While all neurons are modeled as described, we utilize all three compartments only for PYR neurons. For interneurons, we use the basal compartment for all connections and the somatic compartment for neuron dynamics. In biological neurons, the information received by the basal dendrites and apical dendrites is processed in distinct ways. The requirement to use passive dendritic compartments in the model for basal and apical currents is also a result of the circuit implementation: as the activity of the somatic compartment is driven by the *sum* of the basal and apical currents, and the Δ*w* learning circuits rely on their *difference*, the two currents *I*_*a*_ and *I*_*b*_ (eq. (7)) have to be copied into separate pathways and summed in the somatic compartment, and subtracted in the Δ*w* block, as illustrated in figure 1b (top).

In addition to membrane current, neuron also accumulate Calcium trace (*θ*) which is a low-pass filtered of its own spike activity:

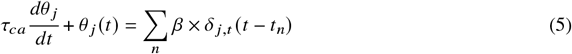

where *β* denotes scaling factor to adjust the contribution of each spike. The neuron’s subthreshold dynamics, the Calcium concentration proxy variable *θ* and error term proportional to (*I*_*a*_ − *I*_*b*_) are all computed in continuous time by the corresponding analog circuit blocks.

### 4.2 Canonical microcircuit attractor dynamics

To achieve the attractor properties of the canonical microcircuit motif, shown in Fig. 1d we modeled two excitatory pools and a shared single inhibitory pool (model parameters are shown in Section SM 3). Following the approach described also in [82], we replaced the recurrent connections within one excitatory population with independent external Poisson spike trains. We then measured the output response of excitatory population to varying input firing rates, mimicking the current flowing within the population though recurrent connections. This allowed us to construct a frequency-frequency (f-f) curve, illustrating the relationship between input and output firing rates in the modified network. Results mirror the reference, revealing two stable fixed points, and the working memory behavior demonstrates sustained activity even after stimulus cessation.

### 4.3 Dendritic currents

The basal and apical dendrites integrate the contribution of multiple synapses. We can express these Excitatory Post-Synaptic Current (EPSC) and Inhibitory Post-Synaptic Current (IPSC) contributions in the following way:

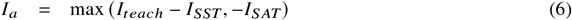

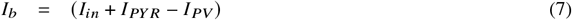

where *I*_*teach*_ is the EPSC produced by the AMPA synapses of the apical dendrite receiving input spikes from the top-down (teacher or prediction) inputs, *I*_*SST*_ represents the IPSC from SST neuron, *I*_*S AT*_ defines the minimum negative plateau current that the apical dendrite circuit can transmit, *I*_*in*_ represents the total weighted sum of all plastic NMDA synapses receiving spikes from the input pathway, *I*_*PY R*_ represents the EPSC produced by the recurrent synapses of the pyramidal cells, and *I*_*PV*_ represents the IPSC produced by the current recurrent inhibition from the PV cells.

Note that in the case of balanced network *I*_*PY R*_ = *I*_*PV*_, and of top-down (attention) stimuli to the VIP neurons *I*_*SST*_ = 0, so the term (*I*_*a*_ − *I*_*b*_) reduces to the classical Delta rule error: (*I*_*teach*_ − *I*_*in*_).

### 4.4 The neuromorphic hardware implementation

The implementation of the somatic compartment, along with the apical and basal dendrites, utilizes existing analog neuromorphic circuits, namely an ultra-low power AdExp-IF neuron circuit [30] and DPI synapses [10, 13]. Apical and basal dendritic compartments are a combination of excitatory and inhibitory DPI synapses, which contribution is summarized into apical (*I*_*a*_) and basal (*I*_*b*_) currents. The sum of *I*_*a*_ and *I*_*b*_ drives the firing activity of the AdExp-IF somatic compartment. This study introduces new circuits to model the Calcium-based dendritic learning rule.

The proposed system includes two independent but complementary learning circuits that drive weight potentiation and depression (Fig. 3a). The upper circuit in Fig. 3a determines the magnitude of weight increase (*I*_*up*_) based on the difference between the basal (*I*_*b*_) and apical (*I*_*a*_) dendritic currents. To prevent unnecessary weight adjustments when the difference between *I*_*b*_ and *I*_*a*_ is minimal, indicating a negligible error, a tolerance term (*I*_*ε*_) is included. This term subtracts from the error, with larger *I*_*ε*_ values increasing the tolerance. The resulting error (*I*_*a*_ - *I*_*b*_ - *I*_*ε*_) is multiplied by a learning rate, adjusting the strength of the weight increase (*I*_*up*_), and is regulated by a third factor voltage variable (*σ*_*LT P*_). If the sum of *I*_*a*_ and *I*_*b*_, integrated in the somatic compartment, sufficiently drives the post-synaptic neuron into the long-term potentiation (LTP) learning region, the final error *I*_*η*_(*I*_*a*_ - *I*_*b*_ - *I*_*ε*_) is processed, leading to an increase in the weight by *I*_*up*_. Conversely, weight depression is managed by a complementary circuit that outputs *I*_*dn*_ when neuron activity is in the long-term depression (LTD) learning region. Utilizing two complementary learning circuits enables better alignment between potentiation and depression, providing the opportunity to apply the same strategies to their layouts.

To harness the benefits of both analog and digital approaches, a novel plastic synapse circuit was designed. This circuit accumulates small errors on a capacitor in the short term, then discretizes and stores weights using standard digital circuitry for long-term retention. This mixed implementation ensures robustness against noise and restricts weight updates to when the accumulated errors on the capacitor are substantial.

Learning signals are continuously computed within the neuron’s somatic block and are available to all plastic synapses. However, individual synapse weight updates occur only upon the arrival of a pre-synaptic spike. When this pre-synaptic input (*V*_*s pk*_) is active, it triggers a positive (*I*_*up*_) or negative current (*I*_*dn*_), which correspondingly increases or decreases the synaptic weight variable *V*_*w*_. In the absence of pre-synaptic input activity, *V*_*w*_ stabilizes towards the middle point between the upper and lower limits (*V*_*dd*_ and ground, respectively) through a Transconductance Amplifier, acting as a stability circuit.

The analog weight value *V*_*w*_ is subsequently digitized and stored using a 4 bit ADC and digital storage circuitry. As *V*_*w*_ reaches its threshold limits, a standard 4 bit digital counter adjusts its counts accordingly, storing these values in Static Random Access Memory (SRAM). Multi-state stability is achieved through the reduced area overhead of digital circuits in advanced technology nodes, such as the 22 nm Fully Depleted Silicon On Insulator (FDSOI) technology used in this work.

## Acknowledgments

We would like to thank Germain Haessig and Dmitrii Zendrikov for initial discussions on some aspects of the learning algorithms presented, Nicolas Brunel and Yiota Poirazi for useful discussions on dendritic processing and cortical computation, Elisabetta Chicca for discussion on the learning circuits and binary classification experiments, and Charl Linssen, Pooja Babu for their support with nest simulator. C.D.L. acknowledges the financial support of the Bridge Fellowship founded by the Digital Society Initiative at University of Zurich (Grant No. G-95017-01-12). This work was also supported by the HORIZON EUROPE EIC Pathfinder Grant ELEGANCE (Grant No. 101161114), and has received funding from Swiss National Science Foundation (SNSF 200021E_222393) and from the Swiss State Secretariat for Education, Research and Innovation (SERI).

## References

[1] G. Indiveri and S.-C. Liu. “Memory and information processing in neuromorphic systems”. In: Proceedings of the IEEE 103.8 (2015), pp. 1379–1397. doi: 10.1109/JPROC.2015.2444094.

[2] S.-C. Liu, T. Delbruck, G. Indiveri, A. Whatley, and R. Douglas. Event-based neuromorphic systems. Wiley, 2014. doi: 10.1002/9781118927601.ch6.

[3] Arindam Basu, Jyotibdha Acharya, Tanay Karnik, Huichu Liu, Hai Li, Jae-Sun Seo, and Chang Song. “Low-Power, Adaptive Neuromorphic Systems: Recent Progress and Future Directions”. In: IEEE Journal on Emerging and Selected Topics in Circuits and Systems 8.1 (Mar. 2018), pp. 6–27. issn: 2156-3365. doi: 10.1109/jetcas.2018.2816339.

[4] C. Mead. “Neuromorphic Electronic Systems”. In: Proceedings of the IEEE 78.10 (1990), pp. 1629–36. doi: 10.1109/5.58356.

[5] R.J. Douglas, M.A. Mahowald, and C. Mead. “Neuromorphic analogue VLSI”. In: Annual Review of Neuroscience 18 (1995), pp. 255–281.

[6] Carver Mead. “Neuromorphic Engineering: In Memory of Misha Mahowald”. In: Neural Computation 35 (2023), pp. 343–383. doi: 10.1162/neco_a_01553.

[7] S.-C. Liu, J. Kramer, G. Indiveri, T. Delbruck, and R.J. Douglas. Analog VLSI: Circuits and Principles. MIT Press, 2002.

[8] M.A. Mahowald. “VLSI analogs of neuronal visual processing: a synthesis of form and function”. PhD thesis. Pasadena, CA.: Department of Computation and Neural Systems, California Institute of Technology, 1992. doi: 10.7907/4bdw-fg34.

[9] C.A. Mead. Analog VLSI and Neural Systems. Reading, MA: Addison-Wesley, 1989.

[10] E. Chicca, F. Stefanini, C. Bartolozzi, and G. Indiveri. “Neuromorphic electronic circuits for building autonomous cognitive systems”. In: Proceedings of the IEEE 102.9 (Sept. 2014), pp. 1367–1388. issn: 0018-9219. doi: 10.1109/JPROC.2014.2313954.

[11] M. Mahowald and R.J. Douglas. “A silicon neuron”. In: Nature 354 (1991), pp. 515–518.

[12] Kwabena Boahen. “Neuromorphic Microchips”. In: Scientific American 292.5 (2005), pp. 56–63. issn: 0036-8733. doi: 10.1038/scientificamerican0505-56.

[13] C. Bartolozzi and G. Indiveri. “Synaptic dynamics in analog VLSI”. In: Neural Computation 19.10 (Oct. 2007), pp. 2581–2603. doi: 10.1162/neco.2007.19.10.2581.

[14] G. Indiveri, B. Linares-Barranco, T.J. Hamilton, A. van Schaik, R. Etienne-Cummings, T. Delbruck, S.-C. Liu, P. Dudek, P. Häfliger, S. Renaud, J. Schemmel, G. Cauwenberghs, J. Arthur, K. Hynna, F. Folowosele, S. Saighi, T. Serrano-Gotarredona, J. Wijekoon, Y. Wang, and K. Boahen. “Neuromorphic silicon neuron circuits”. In: Frontiers in Neuroscience 5 (2011), pp. 1–23. issn: 1662-453X. doi: 10.3389/fnins.2011.00073.

[15] E. Chicca, D. Badoni, V. Dante, M. D’Andreagiovanni, G. Salina, L. Carota, S. Fusi, and P. Del Giudice. “A VLSI recurrent network of integrate–and–fire neurons connected by plastic synapses with long term memory”. In: IEEE Transactions on Neural Networks 14.5 (Sept. 2003), pp. 1297–1307. doi: 10.1109/TNN.2003.816367.

[16] P. Camilleri, M. Giulioni, V. Dante, D. Badoni, G. Indiveri, B. Michaelis, J. Braun, and P. Del Giudice. “A Neuromorphic aVLSI Network Chip with Configurable Plastic Synapses”. In: Hybrid Intelligent Systems, 2007, HIS 2007. (Best paper award). Los Alamitos, CA, USA: IEEE Computer Society, 2007, pp. 296–301. doi: 10.1109/HIS.2007.60.

[17] J. Schemmel, D. Bruderle, A. Grubl, M. Hock, K. Meier, and S. Millner. “A wafer-scale neuromorphic hardware system for large-scale neural modeling”. In: International Symposium on Circuits and Systems, (ISCAS). IEEE. 2010, pp. 1947–1950.

[18] Ning Qiao, Hesham Mostafa, Federico Corradi, Marc Osswald, Fabio Stefanini, Dora Sumislawska, and Giacomo Indiveri. “A reconfigurable on-line learning spiking neuromorphic processor comprising 256 neurons and 128K synapses”. In: Frontiers in neuroscience 9 (2015), p. 141. doi: 10.3389/fnins.2015.00141.

[19] S. Moradi, N. Qiao, F. Stefanini, and G. Indiveri. “A Scalable Multicore Architecture With Heterogeneous Memory Structures for Dynamic Neuromorphic Asynchronous Processors (DYNAPs)”. In: IEEE Transactions on Biomedical Circuits and Systems 12.1 (Feb. 2018), pp. 106–122. doi: 10.1109/TBCAS.2017.2759700.

[20] Georgios Detorakis, Sadique Sheik, Charles Augustine, Somnath Paul, Bruno U Pedroni, Nikil Dutt, Jeffrey Krichmar, Gert Cauwenberghs, and Emre Neftci. “Neural and synaptic array transceiver: A brain-inspired computing framework for embedded learning”. In: Frontiers in neuroscience 12 (2018), p. 583.

[21] Chetan Singh Thakur, Jamal Lottier Molin, Gert Cauwenberghs, Giacomo Indiveri, Kundan Kumar, Ning Qiao, Johannes Schemmel, Runchun Wang, Elisabetta Chicca, Jennifer Olson Hasler, Jae-sun Seo, Shimeng Yu, Yu Cao, André van Schaik, and Ralph Etienne-Cummings. “Large-Scale Neuromorphic Spiking Array Processors: A Quest to Mimic the Brain”. In: Frontiers in Neuroscience 12 (2018), p. 891. doi: 10.3389/fnins.2018.00891.

[22] Ole Richter, Chenxi Wu, Adrian M Whatley, German Köstinger, Carsten Nielsen, Ning Qiao, and Giacomo Indiveri. “DYNAP-SE2: a scalable multi-core dynamic neuromorphic asynchronous spiking neural network processor”. In: Neuromorphic Computing and Engineering 4.1 (Jan. 2024), p. 014003. doi: 10.1088/2634-4386/ad1cd7.

[23] Sung Hyun Jo, Ting Chang, Idongesit Ebong, Bhavitavya B. Bhadviya, Pinaki Mazumder, and Wei Lu. “Nanoscale memristor device as synapse in neuromorphic systems”. In: Nano letters 10.4 (2010), pp. 1297–1301. doi: 10.1021/nl904092h.

[24] G. Indiveri, B. Linares-Barranco, R. Legenstein, G. Deligeorgis, and T. Prodromakis. “Integration of nanoscale memristor synapses in neuromorphic computing architectures”. In: Nanotechnology 24.38 (2013), p. 384010. doi: 10.1088/0957-4484/24/38/384010.

[25] S. Saighi, C. Mayr, B. Linares-Barranco, T. Serrano-Gotarredona, H. Schmidt, G. Lecerf, J. Tomas, J. Grollier, S. Boyn, F. Alibart, S. La Barbera, D. Vuillaume, O. Bichler, C. Gamrat, A. Vincent, and D. Querlioz. “Plasticity in memristive devices”. In: Frontiers in Neuroscience 9.51 (2015), pp. 1–16. doi: 10.3389/fnins.2015.00051.

[26] Z. Wang, S. Joshi, S. E. Savel’ev, H. Jiang, R. Midya, P. Lin, M. Hu, N. Ge, J. P. Strachan, Z. Li, Q. Wu, M. Barnell, G.-L. Li, H. L. Xin, R. S. Williams, Q. Xia, and J. J. Yang. “Memristors with diffusive dynamics as synaptic emulators for neuromorphic computing”. In: Nature materials 16.1 (2017), p. 101.

[27] Suhas Kumar, Xinxin Wang, John Paul Strachan, Yuchao Yang, and Wei D. Lu. “Dynamical memristors for higher-complexity neuromorphic computing”. In: Nature Reviews Materials 7.7 (Apr. 2022), pp. 575–591. issn: 2058-8437. doi: 10.1038/s41578-022-00434-z.

[28] Dennis Valbjørn Christensen et al. “2022 roadmap on neuromorphic computing and engineering”. In: Neuromorphic Computing and Engineering (2022). doi: 10.1088/2634-4386/ac4a83.

[29] S. Ambrogio, P. Narayanan, A. Okazaki, A. Fasoli, C. Mackin, K. Hosokawa, A. Nomura, T. Yasuda, A. Chen, A. Friz, M. Ishii, J. Luquin, Y. Kohda, N. Saulnier, K. Brew, S. Choi, I. Ok, T. Philip, V. Chan, C. Silvestre, I. Ahsan, V. Narayanan, H. Tsai, and G. W. Burr. “An analog-AI chip for energy-efficient speech recognition and transcription”. In: Nature 620.7975 (Aug. 2023), pp. 768–775. issn: 1476-4687. doi: 10.1038/s41586-023-06337-5.

[30] Arianna Rubino, Can Livanelioglu, Ning Qiao, Melika Payvand, and Giacomo Indiveri. “Ultra-Low-Power FDSOI Neural Circuits for Extreme-Edge Neuromorphic Intelligence”. In: IEEE Transactions on Circuits and Systems I: Regular Papers 68.1 (2020), pp. 45–56. doi: 10.1109/TCSI.2020.3035575.

[31] M.J.M. Pelgrom, A.C.J. Duinmaijer, and A.P.G. Welbers. “Matching properties of MOS transistors”. In: IEEE Journal of Solid-State Circuits 24.5 (Oct. 1989), pp. 1433–1440. doi: 10.1109/JSSC.1989.572629.

[32] Wen Sun, Bin Gao, Miaofang Chi, Qiangfei Xia, J Joshua Yang, He Qian, and Huaqiang Wu. “Understanding memristive switching via in situ characterization and device modeling”. In: Nature communications 10.1 (2019), pp. 1–13. doi: 10.1038/s41467-019-11411-6.

[33] Jonathan Timcheck, Jonathan Kadmon, Kwabena Boahen, and Surya Ganguli. “Optimal noise level for coding with tightly balanced networks of spiking neurons in the presence of transmission delays”. In: PLOS Computational Biology 18.10 (Oct. 2022), pp. 1–21. doi: 10.1371/journal.pcbi.1010593.

[34] Thomas SJ Burger, Michael E Rule, and Timothy O’Leary. Active Dendrites Enable Robust Spiking Computations Despite Timing Jitter. bioRxiv:2023.03.22.533815. Mar. 2023. doi: 10.1101/2023.03.22.533815.

[35] N. Perez-Nieves, V.C.H. Leung, P.L. Dragotti, and D.F.M. Goodman. “Neural heterogeneity promotes robust learning”. In: Nature Communications 12 (1 2021), p. 5791. issn: 2041-1723. doi: 10.1038/s41467-021-26022-3.

[36] Jumpei Ukita and Kenichi Ohki. “Adversarial attacks and defenses using feature-space stochasticity”. In: Neural Networks (Aug. 2023). issn: 0893-6080. doi: 10.1016/j.neunet.2023.08.022.

[37] Merav Stern, Nicolae Istrate, and Luca Mazzucato. “A reservoir of timescales emerges in recurrent circuits with heterogeneous neural assemblies”. In: eLife 12 (Dec. 2023). issn: 2050-084X. doi: 10.7554/elife.86552.

[38] Hanle Zheng, Zhong Zheng, Rui Hu, Bo Xiao, Yujie Wu, Fangwen Yu, Xue Liu, Guoqi Li, and Lei Deng. “Temporal dendritic heterogeneity incorporated with spiking neural networks for learning multi-timescale dynamics”. In: Nature Communications 15.1 (Jan. 2024). issn: 2041-1723. doi: 10.1038/s41467-023-44614-z.

[39] Simone D’Agostino, Filippo Moro, Tristan Torchet, Yiğit Demirağ, Laurent Grenouillet, Niccolò Castellani Giacomo Indiveri, Elisa Vianello, and Melika Payvand. “DenRAM: neuromorphic dendritic architecture with RRAM for efficient temporal processing with delays”. In: Nature Communications 15.1 (Apr. 2024). issn: 2041-1723. doi: 10.1038/s41467-024-47764-w.

[40] Dmitrii Zendrikov, Sergio Solinas, and Giacomo Indiveri. “Brain-inspired methods for achieving robust computation in heterogeneous mixed-signal neuromorphic processing systems”. In: Neuromorphic Computing and Engineering 3.3 (July 2023), p. 034002. issn: 2634-4386. doi: 10.1088/2634-4386/ace64c.

[41] Valentina Baruzzi, Giacomo Indiveri, and Silvio P. Sabatini. “Recurrent models of orientation selectivity enable robust early-vision processing in mixed-signal neuromorphic hardware”. In: Nature Communications 16.1 (Jan. 2025). issn: 2041-1723. doi: 10.1038/s41467-024-55749-y.

[42] Yue Kris Wu, Christoph Miehl, and Julijana Gjorgjieva. “Regulation of circuit organization and function through inhibitory synaptic plasticity”. In: Trends in Neurosciences (2022). issn: 0166-2236. doi: 10.1016/j.tins.2022.10.006.

[43] Johannes J Letzkus, Steffen BE Wolff, and Andreas Lüthi. “Disinhibition, a circuit mechanism for associative learning and memory”. In: Neuron 88.2 (2015), pp. 264–276.

[44] Dongchen Liang and Giacomo Indiveri. “Robust state-dependent computation in neuromorphic electronic systems”. In: Biomedical Circuits and Systems Conference (BioCAS). Vol. 2018-Janua. IEEE, Oct. 2018, pp. 1–4. isbn: 978-1-5090-5803-7. doi: 10.1109/BIOCAS.2017.8325075.

[45] Joseph M Brader, Walter Senn, and Stefano Fusi. “Learning real-world stimuli in a neural network with spike-driven synaptic dynamics”. In: Neural Computation 19.11 (2007), pp. 2881–2912.

[46] S. Mitra, S. Fusi, and G. Indiveri. “Real-time classification of complex patterns using spike-based learning in neuromorphic VLSI”. In: Biomedical Circuits and Systems, IEEE Transactions on 3.1 (Feb. 2009), pp. 32–42. doi: 10.1109/TBCAS.2008.2005781.

[47] Lyes Khacef, Philipp Klein, Matteo Cartiglia, Arianna Rubino, Giacomo Indiveri, and Elisabetta Chicca. “Spike-based local synaptic plasticity: a survey of computational models and neuromorphic circuits”. In: Neuromorphic Computing and Engineering 3.4 (Nov. 2023), p. 042001. issn: 2634-4386. doi: 10.1088/2634-4386/ad05da.

[48] Nikos Malakasis, Spyridon Chavlis, and Panayiota Poirazi. “Synaptic turnover promotes efficient learning in bio-realistic spiking neural networks”. In: bioRxiv (2023), p. 2023.05.22.541722. doi: 10.1101/2023.05.22.541722.

[49] Brendan A. Bicknell and Michael Häusser. “A synaptic learning rule for exploiting nonlinear dendritic computation”. In: Neuron 109.24 (Dec. 2021), 4001–4017.e10. issn: 0896-6273. doi: 10.1016/j.neuron.2021.09.044.

[50] Robert Urbanczik and Walter Senn. “Learning by the dendritic prediction of somatic spiking”. In: Neuron 81.3 (2014), pp. 521–528. doi: 10.1016/j.neuron.2013.11.030.

[51] Nicolas Frémaux and Wulfram Gerstner. “Neuromodulated spike-timing-dependent plasticity, and theory of three-factor learning rules”. In: Front. Neur. Circ. 9 (2016), p. 85.

[52] Rodney J. Douglas, Kevan A.C. Martin, and David Whitteridge. “A Canonical Microcircuit for Neocortex”. In: Neural Computation 1 (1989), pp. 480–488. doi: 10.1162/neco.1989.1.4.480.

[53] Guangyu Robert Yang, John D. Murray, and Xiao-Jing Wang. “A dendritic disinhibitory circuit mechanism for pathway-specific gating”. In: Nature Communications 7.1 (Sept. 2016). issn: 2041-1723. doi: 10.1038/ncomms12815.

[54] Charlotte Frenkel, David Bol, and Giacomo Indiveri. “Bottom-Up and Top-Down Approaches for the Design of Neuromorphic Processing Systems: Tradeoffs and Synergies Between Natural and Artificial Intelligence”. In: Proceedings of the IEEE 111.6 (June 2023), pp. 623–652. issn: 1558-2256. doi: 10.1109/jproc.2023.3273520.

[55] Peter Sterling and Simon Laughlin. Principles of neural design. MIT Press, 2015. isbn: 9780262534680.

[56] B. Widrow and M.E. Hoff. “Adaptive Switching Circuits”. In: 1960 IRE WESCON Convention Record, Part 4. New York: IRE, 1960, pp. 96–104.

[57] F. Rosenblatt. “The perceptron: A probabilistic model for information storage and organization in the brain”. In: Psychological Review 65.6 (Nov. 1958), pp. 386–408. doi: 10.1037/h0042519.

[58] E.L. Bienenstock, L.N. Cooper, and P.W. Munro. “Theory for the development of neuron selectivity: orientation specificity and binocular interaction in visual cortex”. In: Jour. Neurosci. 2.1 (1982), pp. 32–48.

[59] G-Q. Bi and M-M. Poo. “Synaptic Modifications in Cultured Hippocampal Neurons: Dependence on Spike Timing, Synaptic Strength, and Postsynaptic Cell Type”. In: Journal of Neuroscience 18.24 (1998), pp. 10464–10472.

[60] C. Clopath and W. Gerstner. “Voltage and spike timing interact in STDP – a unified model”. In: Frontiers in Synaptic Neuroscience 2.25 (2010), pp. 1–11. doi: 10.3389/fnsyn.2010.00025.

[61] H. Markram, W. Gerstner, and P.J. Sjöström. “Spike-timing-dependent plasticity: a comprehensive overview”. In: Frontiers in Synaptic Neuroscience 4.2 (2012), pp. 1–3. doi: 10.3389/fnsyn.2012.00002.

[62] Wulfram Gerstner, Marco Lehmann, Vasiliki Liakoni, Dane Corneil, and Johanni Brea. “Eligibility traces and plasticity on behavioral time scales: experimental support of neohebbian three-factor learning rules”. In: Front. Neur. Circ. 12 (2018), p. 53.

[63] Katie C. Bittner, Aaron D. Milstein, Christine Grienberger, Sandro Romani, and Jeffrey C. Magee. “Behavioral time scale synaptic plasticity underlies CA1 place fields”. In: Science 357.6355 (Sept. 2017), pp. 1033–1036. issn: 1095-9203. doi: 10.1126/science.aan3846.

[64] Big Data Needs a Hardware Revolution. Editorial. Feb. 2018. doi: 10.1038/d41586-018-01683-1.

[65] Adnan Mehonic and Anthony J Kenyon. “Brain-inspired computing: We need a master plan”. In: Nature 604.7905 (2022), pp. 255–260. doi: 10.1038/s41586-021-04362-w.

[66] Anthony Zador, Sean Escola, Blake Richards, Bence Ölveczky, Yoshua Bengio, Kwabena Boahen, Matthew Botvinick, Dmitri Chklovskii, Anne Churchland, Claudia Clopath, James DiCarlo, Surya Ganguli, Jeff Hawkins, Konrad Körding, Alexei Koulakov, Yann LeCun, Timothy Lillicrap, Adam Marblestone, Bruno Olshausen, Alexandre Pouget, Cristina Savin, Terrence Sejnowski, Eero Simoncelli, Sara Solla, David Sussillo, Andreas S. Tolias, and Doris Tsao. “Catalyzing next-generation Artificial Intelligence through NeuroAI”. In: Nature Communications 14.1 (Mar. 2023). issn: 2041-1723. doi: 10.1038/s41467-023-37180-x.

[67] Rodney J. Douglas and Kevan A.C. Martin. “Recurrent neuronal circuits in the neocortex”. In: Current Biology 17.13 (2007), R496–R500. doi: 10.1016/j.cub.2007.04.024.

[68] D.J. Amit and G. Mongillo. “Spike-driven synaptic dynamics generating working memory states”. In: Neural Computation 15.3 (2003), pp. 565–596.

[69] Nicolas Brunel and Xiao-Jing Wang. “What Determines the Frequency of Fast Network Oscillations With Irregular Neural Discharges? I. Synaptic Dynamics and Excitation-Inhibition Balance”. In: Journal of Neurophysiology 90.1 (July 2003), pp. 415–430. issn: 1522-1598. doi: 10.1152/jn.01095.2002.

[70] C. van Vreeswijk and H. Sompolinsky. “Chaotic Balanced State in a Model of Cortical Circuits”. In: Neural Computation 10.6 (Aug. 1998), pp. 1321–1371. issn: 1530-888X. doi: 10.1162/089976698300017214.

[71] M. V. Nair and G. Indiveri. “An Ultra-Low Power Sigma-Delta Neuron Circuit”. In: International Symposium on Circuits and Systems, (ISCAS). May 2019, pp. 1–5. doi: 10.1109/ISCAS.2019.8702500.

[72] Karen Adam, Adam Scholefield, and Martin Vetterli. “Sampling and reconstruction of bandlimited signals with multi-channel time encoding”. In: IEEE Transactions on Signal Processing 68 (2020), pp. 1105–1119. doi: 10.1109/TSP.2020.2967182.

[73] Rachel Sava, Elisa Donati, and Giacomo Indiveri. “Feed-forward and recurrent inhibition for compressing and classifying high dynamic range biosignals in spiking neural network architectures”. In: Biomedical Circuits and Systems Conference (BioCAS). IEEE, Oct. 2023. doi: 10.1109/biocas58349.2023.10388963.

[74] L. Le Cam. “The Central Limit Theorem Around 1935”. In: Statistical Science 1.1 (Feb. 1986). issn: 0883-4237. doi: 10.1214/ss/1177013818.

[75] Sophie Denève and Christian K Machens. “Efficient codes and balanced networks”. In: Nature Neuroscience 19.3 (2016), pp. 375–382. issn: 1097-6256. doi: 10.1038/nn.4243.

[76] P. Livi and G. Indiveri. “A current-mode conductance-based silicon neuron for Address-Event neuromorphic systems”. In: International Symposium on Circuits and Systems, (ISCAS). IEEE. May 2009, pp. 2898–2901. doi: 10.1109/ISCAS.2009.5118408.

[77] C. Rossant, D.F.M. Goodman, J. Platkiewicz, and R. Brette. “Automatic fitting of spiking neuron models to electrophysiological recordings”. In: Frontiers in Neuroinformatics (2010), pp. 1–14. issn: ISSN 1662-5196. doi: 10.3389/neuro.11/002.2010.

[78] Daniel J. Amit. Modeling brain function: The world of attractor neural networks. Cambridge University Press, 1992. isbn: 9780521421249.

[79] J.J. Hopfield and D.W. Tank. “Computing with neural circuits-A model”. In: Science 233.4764 (1986), pp. 625–633.

[80] A. Renart, N. Brunel, and X. Wang. “Mean field theory of irregularly spiking neuronal populations and working memory in recurrent cortical networks”. In: Computational Neuroscience: A Comprehensive Approach. Ed. by J. Feng. Boca Raton: Chapman and Hall, 2003, pp. 431–490.

[81] E. Neftci, J. Binas, U. Rutishauser, E. Chicca, G. Indiveri, and R. Douglas. “Synthesizing cognition in neuromorphic electronic systems”. In: Proceedings of the National Academy of Sciences 110.37 (2013), E3468–E3476.

[82] M. Giulioni, P. Camilleri, M. Mattia, V. Dante, J. Braun, and P. Del Giudice. “Robust working memory in an asynchronously spiking neural network realized in neuromorphic VLSI”. In: Frontiers in Neuroscience 5.149 (2012). issn: 1662-453X. doi: 10.3389/fnins.2011.00149.

[83] Maryada, Saray Soldado-Magraner, Martino Sorbaro, Rodrigo Laje, Dean V. Buonomano, and Giacomo Indiveri. “Stable recurrent dynamics in heterogeneous neuromorphic computing systems using excitatory and inhibitory plasticity”. In: bioRxiv (Aug. 2023). doi: 10.1101/2023.08.14.553298.

[84] Łukasz Kuśmierz, Takuya Isomura, and Taro Toyoizumi. “Learning with three factors: modulating Hebbian plasticity with errors”. In: Current Opinion in Neurobiology 46 (Oct. 2017), pp. 170–177. issn: 0959-4388. doi: 10.1016/j.conb.2017.08.020.

[85] M. Graupner and N. Brunel. “Calcium-based plasticity model explains sensitivity of synaptic changes to spike pattern, rate, and dendritic location”. In: Proceedings of the National Academy of Sciences 109 (2012), pp. 3991–3996. doi: 10.1073/pnas.1109359109.

[86] D.J. Amit and S. Fusi. “Dynamic learning in neural networks with material synapses”. In: Neural Computation 6 (1994), p. 957.

[87] T. Pfeil, T. C. Potjans, S. Schrader, W. Potjans, J. Schemmel, M. Diesmann, and K. Meier. “Is a 4-bit synaptic weight resolution enough? - Constraints on enabling spike-timing dependent plasticity in neuromorphic hardware”. In: Frontiers in Neuroscience 6 (2012). issn: 1662-453X. doi: 10.3389/fnins.2012.00090.

[88] Ning Qiao, Chiara Bartolozzi, and Giacomo Indiveri. “An Ultralow Leakage Synaptic Scaling Homeostatic Plasticity Circuit With Configurable Time Scales up to 100 ks”. In: IEEE Transactions on Biomedical Circuits and Systems (2017). doi: 10.1109/TBCAS.2017.2754383.

