## Supplementary material for "A canonical cortical electronic circuit for neuromorphic intelligence"

### Supplementary material:
### A canonical cortical electronic circuit for neuromorphic intelligence

Maryada[*1], Chiara De Luca[*1,2], Arianna Rubino[3], Chenxi Wen[1], Matteo Cartiglia[4], Ioan-Iustin Fodoruț[5], Melika Payvand[1], and Giacomo Indiveri[1]

[1]Institute of Neuroinformatics, University of Zurich and ETH Zurich, Zurich, Switzerland
[2]Digital Society Initiative, University of Zurich, Zurich, Switzerland
[3]Department of Information Technology and Electrical Engineering, ETH Zurich
[4]IMEC, Leuven, Belgium
[5]Astrali, Romania

#### Supplementary Material

##### SM 0.1 Neuron and synapse model

To create a biologically-inspired SNN model that aligns with a heterogeneous analog hardware implementation, it is imperative to faithfully replicate the circuit dynamics with it's inherent variability. As stated earlier, the chip emulates Adaptive Exponential Integrate-and-Fire (AdExp-IF) neuron model with (alpa/exp) synapse. All synapses are represented with a fixed resolution (3 and 4 bits respectively static and plastic synapses). The somatic membrane current is described as a second order Differential Pair Integrator (DPI) circuit [1] and is given by:

$$\frac{dI_{mem}}{dt} = \frac{\alpha \cdot (I_{in} - I_{sum}) - (I_{sum}) \cdot I_{mem}/I_{dpi,\tau}}{\left(\tau^{soma} \cdot \left(1 + I_{dpi,\tau}/I_{mem}\right)\right)} \tag{1}$$

where

$$I_{sum} = I_{dpi,\tau} - I_{pfb} + I^{basal}_{gaba_a}, \tag{2}$$

Positive feedback current ,

$$I_{pbf} = k_{pbf} \cdot \left(I_{mem} - p_{pfb} \cdot I_{pfb,th}\right) \tag{3}$$

Adaptation current is:

$$\frac{dI_{ahp}}{dt} = -\frac{1}{\tau_{ahp}} \cdot I_{ahp}, \tag{4}$$

$$\text{if spike: } I_{ahp} \mathrel{+}= I_{ahp,w0} \cdot \left(\alpha_{ahp}\right)^{\frac{k_n}{k_p}} \cdot \left(\frac{I_\tau}{I_{ahp}}\right)^{\frac{k_n}{k_p}-1} \tag{5}$$

$$\tag{6}$$

The incoming basal currents are defined as:

$$I_{in} = I_{const} - I_{ahp} + I^{apical} + I^{basal}_{ampa} + \hat{I}^{basal}_{nmda} - I^{basal}_{gaba_b} \tag{7}$$

where $\hat{I}^{basal}_{nmda}$ is voltage-dependent receptor,

$$I^{basal}_{nmda} < I_{nmda,th} \; ? \; \hat{I}^{basal}_{nmda} = I^{basal}_{nmda} : \hat{I}^{basal}_{nmda} = 0 \tag{8}$$

*These authors have contributed equally to this work.

Whenever a somatic spike is emitted ($I_{mem}^{soma} > I_{th}^{soma}$), the neuron will stay silent for a refractory period r, the membrane current $I_{mem}^{soma}$ will be reset at a fixed $I_{reset}^{soma}$ value.

The second order DPI for qll receptors ($N$-Methyl-D-aspartate (NMDA), $\alpha$-Amino-3-hydroxy-5-methyl-4-isoxazolepropionic Acid (AMPA), $\gamma$-Aminobutanoic Acid (GABA)_A, GABA_A) and Calcium dynamics is given by:

$$\frac{dI_w}{dt} = -\frac{-I_w}{\tau};$$
$$\text{if spike: } I_w + = \alpha \cdot I_{w0}$$
$$\frac{dI}{dt} = \frac{1}{\tau} \cdot \left( -I + \frac{I_w}{I_\tau} \cdot \frac{I}{1 + I/I_g} \right)$$

To enable variability in all neural and synapse dynamics, each parameter $p$ is independently drawn from a uniform distribution: $[p - \sigma p, p + \sigma p]$ where $\sigma$ can be defined by the user.

**Apical dendritic compartment:** The apical input $I^{apical}$ is processed by half-wave rectifier circuit and is defined as follows:

$$I^{apical} = \begin{cases} I_{ampa}^{apical} - I_{gaba_b}^{apical} & \text{if} \quad I_{ampa}^{apical} - I_{gaba_b}^{apical} > I_{th}^{apical} \\ I_{th}^{apical} & \text{otherwise} \end{cases} \tag{9}$$

#### SM 0.2 Comparison of circuit simulation and model

To validate the neuron and synapse models, we compare the traces recorded from NEST simulation with detailed circuit-level simulations. In Figure S1, we show the neuron membrane current (see Fig. S1a), adaptation current (see Fig. S1b) when a regular stimulus (see Fig. S1f)is inject to somatic compartment for both circuit simulation and model simulation. The behaviour of both the positive feedback mechanism (see Fig. S1c) and the second-order DPI calcium current integrator (see Fig. S1d,S1e) are also shown. The mismatch observed in the former is due to temporal resolution of simulation, not always able to capture the injected current from this dynamics. The behaviour of the receptor dynamics implemented in chip as a second order DPI is also consistent between behavioural and circuit simulation: specifically we compared the neuronal receptor dynamics when stimulated by a single receptor with regular spike train. It is shown an almost perfect overlap between circuit and model simulation for both first order and second order receptor dynamics. This representative simulation, we demonstrate an optimal scenario where the dynamics of key variables in the NEST behavioral simulator align with those of the circuit simulation implemented in Cadence. Although the biases in real silicon may not be precise enough to perfectly match the NEST simulation, the simulation successfully validates the expected circuit dynamics

### SM 1 BCM curve

The Bienenstock-Cooper-Munro (BMC) theory [3] posits that a synapse strengthens when the pre- and post-synaptic neurons fire together and the average activity of the neuron exceeds a certain threshold, with the threshold itself adapting based on the neuron's past activity.

This work presents a third-factor learning rule, where the third factor, i.e. the calcium concentration of the post-synaptic neuron, acts as a binary signal to enable learning. The magnitude of weight change follows the Delta rule as described in Section 2.1. We analyzed the trends in weight changes to compare the dynamics of this learning rule with those of BCM theory. Figure S2 illustrates the relationship between synaptic weight change ($\Delta w$) and post-synaptic neuron calcium current ($\theta$).

Figure S2 illustrates the relationship between Long Term Potentiation (LTP) and Long Term Depression (LTD) learning regions with respect to post-synaptic calcium activity. The vertical dashed lines indicate the lower and upper learning thresholds, highlighted by green (LTD) and red (LTP) horizontal lines. The shaded gray area represents the standard deviation, reflecting variability in these regions. This plot demonstrates the stop-learning mechanism for both low and high values of calcium current, while clearly distinguishing the LTD and LTP regions.

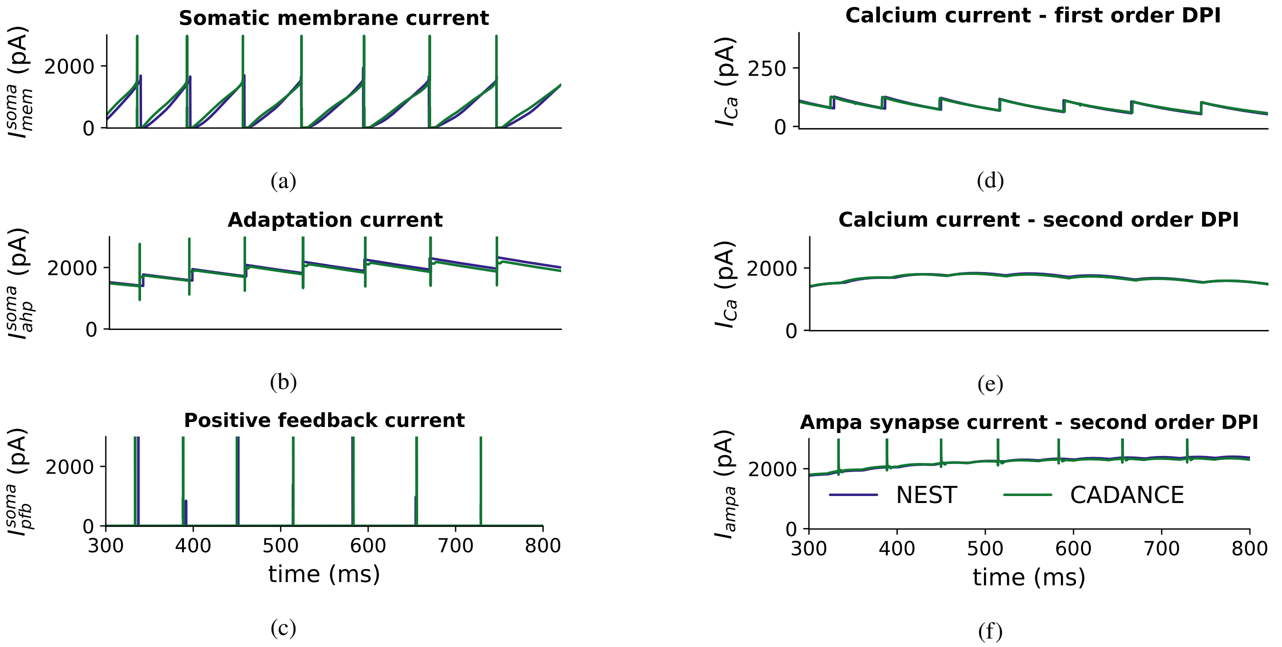


Figure S1: Neuron response to a regular 25 Hz input spike train through an AMPA synapse. High-level software simulations using the NEST simulator [2] are shown in blue, and low-level electronic circuit simulations in green. (a) somatic membrane current; (b) adaptation current; (c) positive feedback current; (d) $1^{st}$ order DPI filter output of the somatic membrane current; (e) $2^{ns}$ order output; (f) AMPA synapse output current.

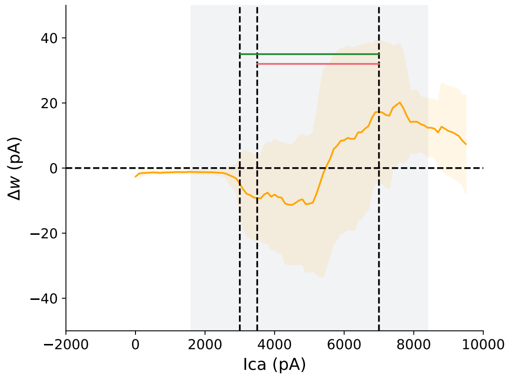


Figure S2: Rolling average of the weight update distribution versus the postsynaptic calcium current during the training phase of a classification task (same experiment performed in the main text, section 2.1). Vertical dotted lines depict the calcium LTP and LTD regions, also highlighted by the green and red horizontal lines on top respectively. Gray area shows the interval of confidence of such thresholds when affected by a 20% device mismatch variability.

### SM 2 Learning and simulation paradigm

This section delves into the specifics of the training-testing protocols and pre-processing methods employed during simulations. First, we validate our model by learning two patterns with varying degrees of overlap. As shown in figure 2c in the main text, the network can distinguish between two patterns with an accuracy rate exceeding 80%, even with a 90% overlap between them.

#### SM 2.1 Binary classification

In this experiment, the network is trained to classify binary patterns of size 64 using a supervised learning approach. These patterns are composed of binary vectors from independent Poisson spike trains, exhibiting either a high mean firing rate (50 Hz) or a low one (5 Hz). Each pattern is created with $0.5 \times (1 + 0.5 * 0.5 * ov) \times n_{inp}$ high rates and $0.5 \times (1 - 0.5 * 0.5 * ov) \times n_{inp}$ low-rate elements, where $ov$ denotes the overlap ratio. (i.e.

| | |
|---|---|
| Presentation duration | 1 sec |
| Seed range | [1-20] |
| Number of Repetitions | 20 |
| Sampled presented | ... |

Table S1: Simulation parameters

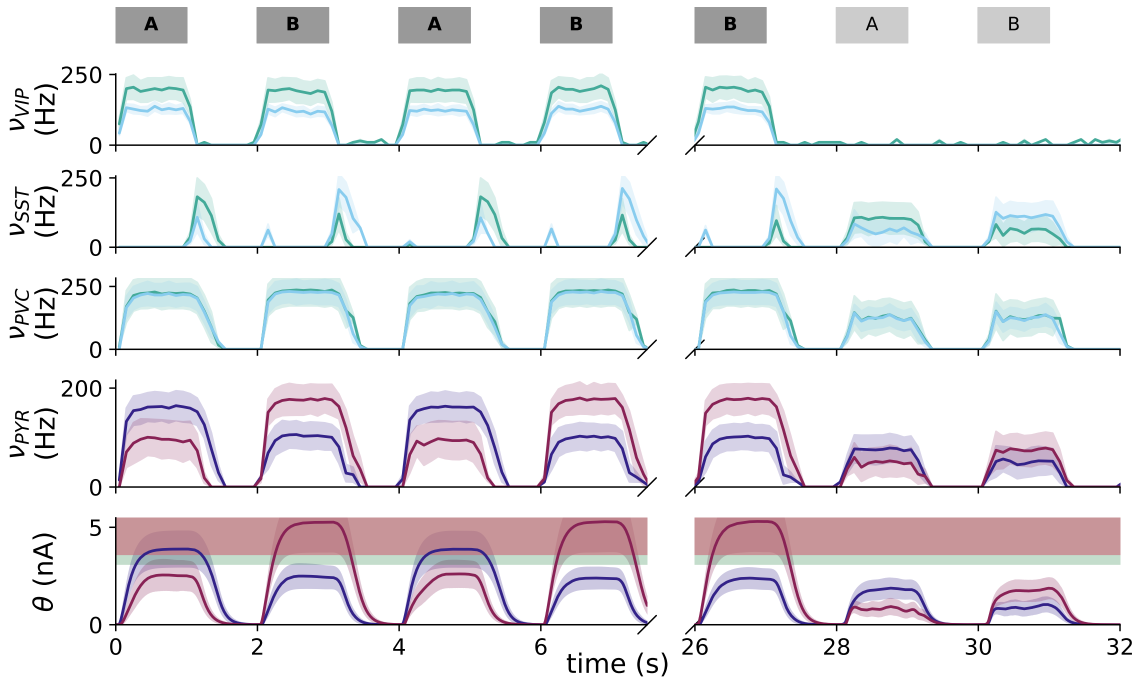


Figure S3: Network activity for an orthogonal (non-overlapping) pattern experiment

synapses characterized by the same high status whenever both patterns are presented).

During training, a localized high frequency teacher signal (Poisson rate = 50 Hz) is applied as a top-down apical input to the pyramidal neurons of a designated column. Simultaneously, Vasoactive Intestinal Peptide (VIP) neurons in all columns receive a high-frequency (50 Hz) "contextual" signal though AMPA synapses, targeting the basal compartment. Note that all interneurons are modeled as point neurons.

During the testing phase, a low teacher signal (rate = 5 Hz) is applied as apical input to all pyramidal neurons and VIP neurons in the mico-circuit, simulating noise from other brain areas. This signal is insufficient to strongly activate VIP cells, allowing Somatostatin (SST) inhibitory cells to remain active due to Pyramidal (PYR) projections. Additionally, a negative current is injected into the pyramidal population through the apical compartment, following the half-wave rectification described in SM 0.1. This mechanism is crucial for maintaining low firing rates in pyramidal neurons, limiting calcium dynamics, and preventing synaptic weight change under noisy conditions mimicking real-world scenarios. This behavior, which can be easily fine-tuned, ensures a halt in learning and prevents the forgetting of memories in noisy settings. Figure S3 illustrate the activity of neuron population for orthogonal input pattern. We assessed the network performance across a range of overlaps with varying degrees of device mismatch. Figure S4a shows performance accuracy over epoch for a fixed overlap of 50%. We further analyzed the performance with various degrees of overlap and device mismatch, figure S4b shows decline in accuracy with increased device mismatch.

This columnar organization for binary classification can be easily extended to multi-class classification by using multiple local "columnar" copies of the circuit, with local lateral inter-columnar connectivity: nearby populations share inputs to their local Parvalbumin (PV) cell clusters, to implement a Winner-Take-All (WTA) functionality and enhance the separability of the responses to the different classes being trained. From the hardware implementation point of view, the hierarchical multi-core approach used in many mixed-signal

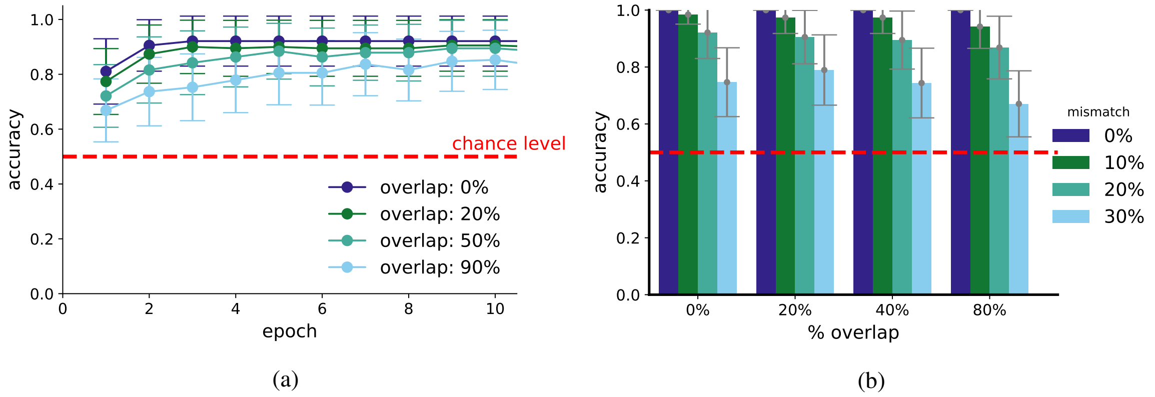


Figure S4: Effect of different mismatch and overlap percentages in the accuracy of the binary classification experiment of Section2.2. (a) Accuracy versus the amount of overlap in the input pattern, for a conservative mismatch variability figure of 20%. (b) Plateau accuracy for various mismatch and overlaps. Statistics is acquired over 20 independent runs.

neuromorphic processors [4–6] can directly support the small-world network connectivity patterns of this columnar approach.

##### SM 2.1.1 Functional consequences of inhibitory neuron disruption

Section 2.1 explains the functioning of the canonical microcircuit and the role of each neuron during different phases of the learning and inference processes. To validate this, lesion (i.e., knockout) studies were conducted on PV and SST neurons. For PV, we severed the connections from PV to PYR cells during both phases. PV cells facilitates WTA mechanism and maintain balanced activity in a recurrent network. In the absence of inhibition via PV, all PYR neurons in both the clusters saturates to their highest firing rate (see fig. S5a), confirming that these functionalities are indeed mediated by PV cells.

On the other hand, disrupting the connections between SST and pyramidal cells (see figure S5b) had no discernible effect during the learning phase, as expected, since SST cells are inhibited by VIP cells during this phase. However, it did affect the inference phase. The absence of apical inhibition resulted in increased activity during the inference phase, leading to alterations in synaptic weights (see figure S5c).

#### SM 3 Network Parameters

In this paragraph the parameters to reproduce results are listed. In table S2, the simulation parameter characterizing the 6 implemented synaptic receptors are listed. For the sake of simplicity, the derived time constants ($\tau = CU_T/(kI_{tau})$) are summarized in Table S4

#### References

[1] E. Chicca, F. Stefanini, C. Bartolozzi, and G. Indiveri. "Neuromorphic electronic circuits for building autonomous cognitive systems". In: *Proceedings of the IEEE* 102.9 (Sept. 2014), pp. 1367–1388. ISSN: 0018-9219. DOI: 10.1109/JPROC.2014.2313954.

[2] *NESTML: a modeling language for spiking neurons*. Zenodo, Mar. 2016. DOI: 10.5281/zenodo.1412345.

[3] Leon N Cooper. *Theory of cortical plasticity*. World Scientific, 2004.

[4] S. Moradi, N. Qiao, F. Stefanini, and G. Indiveri. "A Scalable Multicore Architecture With Heterogeneous Memory Structures for Dynamic Neuromorphic Asynchronous Processors (DYNAPs)". In: *IEEE Transactions on Biomedical Circuits and Systems* 12.1 (Feb. 2018), pp. 106–122. DOI: 10.1109/TBCAS.2017.2759700.

Table S2: Receptor parameters, all values listed here are affected by mismatch.

| **Receptor type** | **Name** | **Value** | **Description** |
|---|---|---|---|
| Basal AMPA | $\alpha$ | 4 | synpase scaling factor |
| | $I_\tau$ | 5.4 pA | Synaptic time constant current, inversely proportional to $\tau$ |
| Basal NMDA | $\alpha$ | 4 | synpase scaling factor |
| | $I_\tau$ | 0.1 pA | Synaptic time constant current, inversely proportional to $\tau$ |
| | $V_\theta$ | 0 pA | NMDA voltage gating threshold |
| Basal GABA-A | $\alpha$ | 4 | synpase scaling factor |
| | $I_\tau$ | 18.5 pA | Synaptic time constant current, inversely proportional to $\tau$ |
| Basal GABA-B | $\alpha$ | 4 | synpase scaling factor |
| | $I_\tau$ | 12 pA | Synaptic time constant current, inversely proportional to $\tau$ |
| Apical AMPA | $\alpha$ | 4 | synpase scaling factor |
| | $I_\tau$ | 0.46 pA | Synaptic time constant current, inversely proportional to $\tau$ |
| Apical GABA-B | $\alpha$ | 4 | synpase scaling factor |
| | $I_\tau$ | 0.5 pA | Synaptic time constant current, inversely proportional to $\tau$ |

| Neuron parameters | | | |
|---|---|---|---|
| | name | value | description |
| Shared neuron parameters | $C^{mem}_{soma}$ | $1.1815pF$ | membrane capacitance, fixed at layout time |
| | $^*I^{reset}_{soma}$ | $0.05pA$ | post-spike reset value |
| | $^*I^{const}_{soma}$ | $0.pA$ | external constant current |
| | $^*\alpha^{mem}_{ahp}$ | 2 | synpase scaling factor |
| | $C^{mem}_{ahp}$ | $1.1520pF$ | adaptation capacitance, fixed at layout time |
| | $^*I^{soma}_{ahp,tau}$ | $0.2pA$ | adaptation time current, inversely proportional to $\tau$ |
| | $^*I^{soma}_{ahp,bias}$ | $100nA$ | adaptation current bias |
| | $^*\alpha^{mem}_{Ca}$ | 4 | synpase scaling factor |
| | $C^{mem}_{ca}$ | $1.164pF$ | caclium capacitance, fixed at layout time |
| | $^*I^{soma}_{ca,tau}$ | $0.6pA$ | caclium time current, inversely proportional to $\tau$ |
| | $^*I^{soma}_{ca,bias}$ | $300nA$ | caclium current bias |
| | $^*I^{soma}_{pfb,gain}$ | $4000pA$ | positive feedback gain current |
| | $^*I^{soma}_{pfb,norm}$ | $20pA$ | positive feedback normalization current |
| | $^*I^{soma}_{pfb,th}$ | 0.75 | positive feedback threshold ratio |
| pyr neurons | $^*I^{dpi}_{soma,tau}$ | $1.9pA$ | synaptic time constant current, inversely proportional to $I_{tau}$ |
| | $^*I^{dpi}_{soma,th}$ | $4000pA$ | spike threshold current |
| | $^*t_{ref}$ | $5ms$ | spike threshold current |
| PV neurons | $^*I^{dpi}_{soma,tau}$ | $4pA$ | synaptic time constant current, inversely proportional to $I_{tau}$ |
| | $^*I^{dpi}_{soma,th}$ | $9000pA$ | spike threshold current |
| | $^*t_{ref}$ | $2.5ms$ | spike threshold current |
| SST neurons | $^*I^{dpi}_{soma,tau}$ | $2.5pA$ | synaptic time constant current, inversely proportional to $I_{tau}$ |
| | $^*I^{dpi}_{soma,th}$ | $7000pA$ | spike threshold current |
| | $^*t_{ref}$ | $2.5ms$ | spike threshold current |
| VIP neurons | $^*I^{dpi}_{soma,tau}$ | $3.8pA$ | synaptic time constant current, inversely proportional to $I_{tau}$ |
| | $^*I^{dpi}_{soma,th}$ | $11000pA$ | spike threshold current |
| | $^*t_{ref}$ | $2.5ms$ | spike threshold current |

Table S3: Neuron parameters, values marked with $*$ are affected by mismatch.

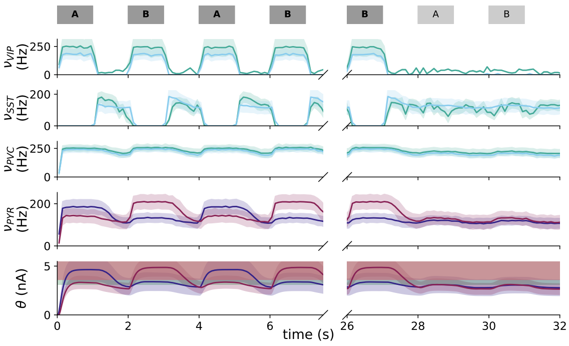


(a) PV knockout experiment

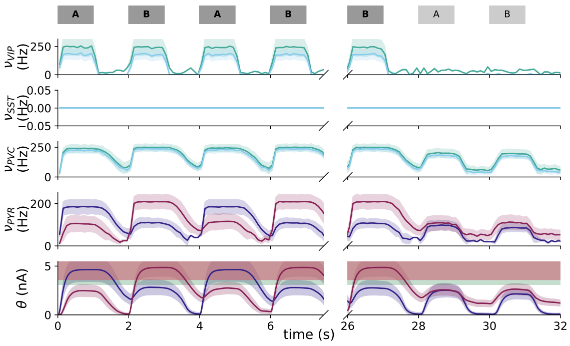


(b) SST knockout experiment

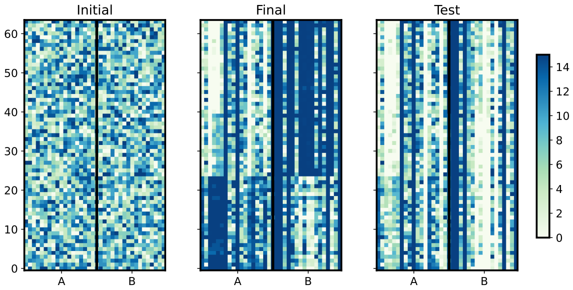


(c)

Figure S5: The experiment described in the main text, section 2.2 has been reproduced by performing knockout experiments on PV and SST inhibitory cells, respectively. (a) PV cells knockout experiment. This experiment explores network activity when PYR → PV connections are absent. In the absence of the winner-take-all inhibition mechanism, no learning takes place. (b): SST cells knockout experiment. This experiment explores network activity when PYR → SST connections are absent. The SST knockout does not affect the learning phase. In both the inference phase and when no input is provided (pause phases), on the other hand, it results in an higher activity of pyramidal cells, leading to catastrophic forgetting. (c) shows the synaptic metrix before the training phase, after the training phase and after the inference phase

| Network time constants | |
|---|---|
| $\tau$ | value (ms) |
| $\tau_{soma}$ | 22 |
| $\tau_{ahp}$ | 500 |
| $\tau_{Ca}$ | 72 |
| $\tau_{ampa}^{basal}$ | 7 |
| $\tau_{ampa}^{apical}$ | 89 |
| $\tau_{nmda}^{basal}$ | 88 |
| $\tau_{gabaA}^{basal}$ | 4 |
| $\tau_{gabaB}^{basal}$ | 4 |
| $\tau_{gabaB}^{apical}$ | 72 |

Table S4: Network time constants

[5] Nishant Mysore, Gopabandhu Hota, Stephen R. Deiss, Bruno U. Pedroni, and Gert Cauwenberghs. "Hierarchical Network Connectivity and Partitioning for Reconfigurable Large-Scale Neuromorphic Systems". In: *Frontiers in Neuroscience* 15 (2022). ISSN: 1662-453X. DOI: 10.3389/fnins.2021.797654.

[6] Ole Richter, Chenxi Wu, Adrian M Whatley, German Köstinger, Carsten Nielsen, Ning Qiao, and Giacomo Indiveri. "DYNAP-SE2: a scalable multi-core dynamic neuromorphic asynchronous spiking neural network processor". In: *Neuromorphic Computing and Engineering* 4.1 (Jan. 2024), p. 014003. DOI: 10.1088/2634-4386/ad1cd7.

| Number of neurons per column | |
|---|---|
| population | value |
| pyr | 18 |
| PV | 5 |
| SST | 5 |
| VIP | 4 |
| input | 64 |
| teacher | 4 |
| context | 4 |

Table S5: Number of neurons per column

| **Column parameters** | | | | | |
|---|---|---|---|---|---|
| | Pyr | PV | SST | VIP | receptor type |
| Pyr | 6\|1.0 | 8 \| 1.0 | 8 \| 1.0 | - | $AMPA$ basal |
| PV | 8\|1.0 | 4 \| 1.0 | - | - | $GABA_B$ basal |
| SST | 8\|1.0 | - | - | - | $GABA_B$ apical |
| VIP | - | - | 8 \| 1.0 | - | $GABA_A$ |
| Ibias | 4000 pA | | | | |

Table S6: Column parameters

| **Inter-column parameters** | | | | | |
|---|---|---|---|---|---|
| | Pyr | PV | SST | VIP | receptor type |
| Pyr | 2 \| 0.5 | 8 \| 0.5 | - | - | $AMPA$ basal |
| Context | - | - | - | 4 \| 1.0 | $AMPA$ basal |
| Teacher | 8 \| 1.0 | - | - | - | $AMPA$ apical |
| Input | random \| 1.0 | 1 \| 1.0 | - | - | $NMDA$ |
| Ibias | 8000 pA | | | | |
| Ibias plastic | 1000 pA | | | | |

Table S7: Inter-column parameters
